# Desolvation Energy Explains Partitioning of Client Proteins into Condensates

**DOI:** 10.1101/2021.08.16.456554

**Authors:** José A. Villegas, Emmanuel D. Levy

## Abstract

Membraneless organelles are cellular compartments that form by liquid-liquid phase separation of one or more components. Other molecules, such as other proteins and nucleic acids, will distribute between the cytoplasm and the liquid compartment in accordance with the thermodynamic drive to lower the free energy of the system. The resulting distribution colocalizes molecular species, to carry out a diversity of functions. Two factors could drive this partitioning: the difference in solvation between the dilute versus dense phase, and intermolecular interactions between the client and scaffold proteins. Here, we develop a set of knowledge-based potentials that allow for the direct comparison between desolvation energy and pairwise interaction energy terms, and use these to examine experimental data from two systems: protein cargo dissolving within phase-separated droplets made from FG repeat proteins of the nuclear pore complex, and client proteins dissolving within phase-separated FUS droplets. We find close agreement between desolvation energies of the client proteins and the experimentally determined values of the partition coefficients, while pairwise interaction energies between client and scaffold show weaker correlations. These results show that client stickiness is sufficient to explain differential partitioning of clients within these two phase-separated systems without taking into account the composition of the condensate. This suggests that selective trafficking of client proteins to distinct membraneless organelles requires recognition elements beyond the client sequence composition.

## INTRODUCTION

Cellular functions require the spatial and temporal organization of a vast number of molecular components. Cells achieve such organization by complex expression programs,(Alberts, 2017) the use of membrane-bound compartments,(Alberts, 2017) chemical gradients,(Howard, 2012; Panbianco and Gotta, 2011) and membrane-less organelles that form through a process of phase separation.(Banani et al., 2017; Brangwynne et al., 2009, 2011; Ge et al., 2009; Li et al., 2012; Zwicker et al., 2014) Membrane-less organelles maintain chemical heterogeneity in the cell by exploiting the differences of solubility of nucleic acids, organic molecules, and other proteins in the aqueous and proteinaceous/nucleic acid phases.(Banani et al., 2017) For example, liquid droplets formed by the RNA-binding protein fused in sarcoma (FUS) can recruit client RNA binding proteins through intermolecular interactions between low complexity (LC) domains.(Kato et al., 2012) Client partitioning within phase-separated liquid droplets can be used together with other cellular strategies for compartmentalization, as in the distribution of molecular species between the cytosol and the nucleus. While the nuclear envelope serves as a barrier to prevent the contents of the cytoplasm and the nucleus from mixing, transport through the nucleus double membrane is mediated by a liquid protein phase composed of intrinsically disordered domains of the nuclear pore complex (NPC). These disordered regions are rich in phenylalanine and glycine residues, and are known as FG domains. Protein cargo must dissolve within the liquid protein phase to gain passage through the pore and entry into the nucleus.(Hülsmann et al., 2012; Ribbeck and Görlich, 2001) In these and other systems, the composition and sequence of a protein’s polypeptide chain determines the chemical potential for going from the aqueous phase to the protein-rich phase. (Frey et al., 2018; Nott et al., 2016; Wang et al., 2018)

Proteins that undergo liquid-liquid phase separation often show extensive disordered regions, and as such undergo complex dynamics (Das and Pappu, 2013; Sawle and Ghosh, 2015) which makes it difficult to characterize the distribution of amino acid pairwise interactions in protein droplets.(Dignon et al., 2018) Insights have been gained through theory, sequence analysis, variation of solution conditions, and mutational analysis. (Dignon et al., 2018; Lin et al., 2017, 2016; Nott et al., 2015; Pak et al., 2016; Regy et al., 2020; Schuster et al., 2020; Vernon and Forman-Kay, 2019; Vernon et al., 2018; Zaslavsky and Uversky, 2018) Early work on the sequence determinants of phase-separation established aromatic interactions as important for driving phase-separation of proteins. Nott *et. al*. found that mutating phenylalanine residues of the intrinsically disordered protein Ddx4 dramatically increase the threshold concentration required for phase-separation.(Nott et al., 2015) Similarly, Lin *et. al*. found that tyrosine residues are critical for the phase-separation behavior of FUS.(Lin et al., 2017) These observations are consistent with π-π interactions being an important attribute of amino-acid interactions. Indeed, Vernon *et. al*. relied on frequencies of amino acids with *sp*^2^ character to train a sequence-based predictor of phase-separation behavior. (Vernon et al., 2018) However, descriptions of phase-separating systems can be constructed without incorporating specific enthalpic interactions. Hardenberg *et al*. used amino acid specific disorder propensities in bound and unbound states to construct a predictive model of phase-separation propensities, where only a non-specific enthalpy term was incorporated to account for hydrophobic interactions.(Hardenberg et al., 2020)

A related but distinct problem is the partitioning of client proteins within phase-separated systems.(Ditlev et al., 2018; Lin et al., 2015; Nott et al., 2016) Although it is increasingly recognized that the phase-separated state is one that is accessible to proteins in general,(Fuxreiter and Vendruscolo, 2021) client proteins are regarded as not constituting an essential component in condensate formation by a phase-separating species. The client protein can however form interactions with the condensed phase and occupy the interior. In such a case, the free energy of interaction between two components (client and condensate) is directly measurable by calculating the ratio of concentrations of the client in the protein-rich phase and the aqueous phases. Prediction of client partitioning within phase-separated systems has been treated theoretically with Monte Carlo simulations that perform particle transfer from the dilute to the dense phase.(Ghosh and Mazarakos, 2019; Qin and Zhou, 2016)

The partition coefficient, or the ratio of concentrations of species dissolved in two phases at equilibrium, is a measure of the change in the free energy for a species going from one phase to another. Transfer energies have been extensively studied for individual amino acids, from an aqueous to a non-aqueous environment, resulting in numerous hydrophobicity and hydropathy scales.(Kyte and Doolittle, 1982; Roseman, 1988; Simm et al., 2016; Wimley and White, 1996) One such scale is residue interface propensity, which provides a statistical estimate of transfer free energies of amino acids from solvent to a protein interface of average composition by comparing the frequency of amino acids at the protein surface *versus* interface.(Levy et al., 2012) In contrast, amino acid pairwise contact preferences have been estimated from over- or under-representation of contacts in the protein interior,(Miyazawa and Jernigan, 1985) or at protein interfaces.(Glaser et al., 2001) It was previously observed that a weighted combination of residue interface propensity and an amino acid interaction potentially was better at distinguishing true protein-protein interfaces from decoys.(Nadalin et al., 2018) Importantly, contact counts (that estimate interaction propensity) and frequency counts (that estimate desolvation energy) were weighted since the two are different quantities that cannot be compared directly (Figure 1).

**Figure 1.**
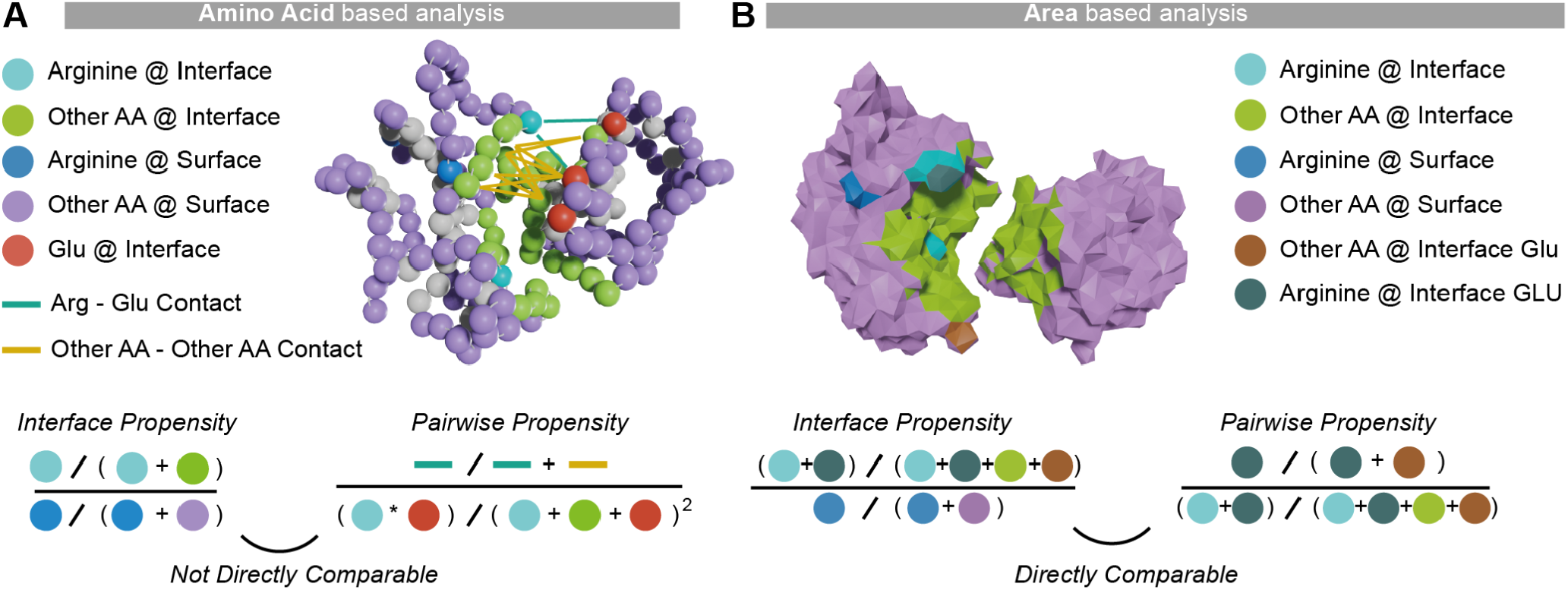
Considering the surface area of amino acids allows a direct comparison of interface propensities and pairwise contact propensities. **A**. The *interface propensity* of arginine is the ratio of arginine frequency at the interface relative to its frequency at the surface. The *pairwise contact propensity* between arginine and glutamate is the frequency of their contacts at the interface relative to their frequency at the interface. These two propensities are not directly comparable because the terms used in their derivation are in different units (amino acid frequencies for the former, and normalized contact frequencies for the latter). **B**. Considering the surface area of amino acids makes interface propensity and contact propensity directly comparable because the same measure (i.e., area fraction) is used to derive all terms. The *interface propensity* of arginine becomes its fractional area at the interface relative to the surface, and the arginine-glutamate *contact propensity* becomes the fractional area of arginine-glutamate contacts relative to the fractional interface area occupied by arginine.

We show here that the desolvation and interaction energies derived from amino acid surface areas are directly comparable without arbitrary weighting. To unify these two descriptions into a single energy term, we compared the same quantity - contact surface area - of amino acids at solvated surfaces versus interfaces. The capability of Voronoi tessellation(Richards, 1974; Singh et al., 1996) to make both descriptions comparable is illustrated in Figure 1. Voronoi tessellation enables the exact subdivision of any protein surface, making it possible to estimate both desolvation energy and contact preferences from fractional amino acid surface areas. We use the derived potentials to show that the partitioning of client proteins within condensates is best explained by the desolvation energy of the client protein. In contrast, we find that pairwise-amino acid interactions between client and scaffold explained the degree of partitioning less well.

## METHODS

### Dataset

The 3DComplex database (Levy et al., 2006) was used to select 1011 heteromeric dimers from a non-redundant set of proteins. The dataset was divided randomly into three sets, analyses were carried out on each set independently. The resulting scales were the average of the three analyses, which also gave the standard deviation of each propensity value. The dataset consisted of structures with a resolution better than 3.0 Å and was non-redundant at a sequence identity level of 70% as defined in 3DComplex. In order to minimize the number of incorrect biological assemblies(Dey and Levy, 2018), we filtered out complexes with a QSbio(Dey et al., 2018) error probability greater than 10%. Voronoi surfaces and contact areas were computed on the first chain in the biological assembly, using the command line program CAD-score.(Olechnovič and Venclovas, 2014) To derive amino acid propensities, we selected surface and interface residues involving significant contact surface area of their side-chain with either the solvent or a protein partner. Selected interface residues had to satisfy two criteria: (i) expose over 25% of their surface area in the monomeric state, and (ii) 50% of that exposed area had to be buried in the complex. Surface residue also had to satisfy two criteria to be included in the analyses: (i) over 25% of their side-chain area was exposed to the solvent, and (ii) no surface area was involved at an interface.

### Definition of the propensities

The absolute residue interface propensity scale is calculated as

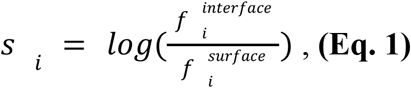

where *s*_*i*_ is the interface propensity of amino acid type *i*, 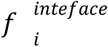 is the area fraction of amino acid type *i* at the interface, and 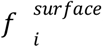 is the area fraction of amino acid type *i* at the solvent-exposed surface. These propensities capture the tendency of amino acids to interact with protein surfaces in general, and as such we also refer to this propensity scale as “stickiness”.(Levy et al., 2012)

The area fraction of amino acid type *i* at the interface is computed as

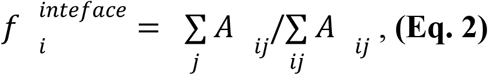

where 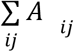 is the total interface area and 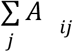 is the surface area of amino acid at the interface.

The area fraction of an amino acid at the surface is obtained as

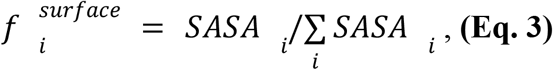

where *SASA*_*i*_ is the total surface area of residues of amino acid type *i* on the first chain in contact with water, as determined by the Voronoi cell capping algorithm.(Olechnovič and Venclovas, 2014) These expressions take the familiar forms used originally to estimate interface propensity scales. (Jones and Thornton, 1996)

The use of the Voronoi tessellation allows us to decompose the interface surface area into residue-level contributions unambiguously. In this manner, we consider an interface as being composed of 20 different sub-interfaces, where each sub-interface corresponds to a single amino acid type. The sub-interface propensity of each amino acid can be calculated as:

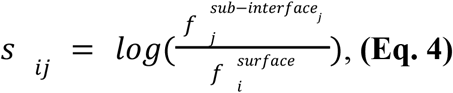

We can re-write this expression as

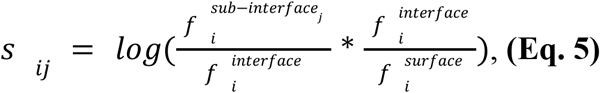

resulting in

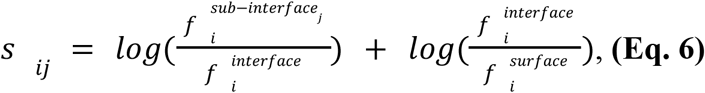

This is equivalent to decomposing the expression into the interaction energy and the desolvation energy, so that we can consider

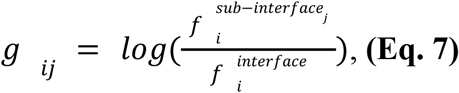

as the energetic contribution of amino acid *i* interacting with the subsurface *j*.

To get the full interaction energy of an amino acid pair, we sum

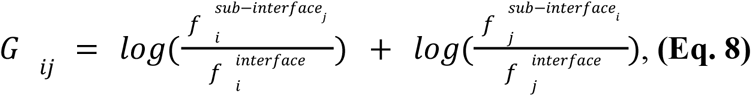

These terms can be combined to yield probability of finding amino acid *i* and amino acid *j* together at the Interface 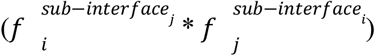, which we can denote as 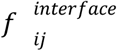. This yields the expression

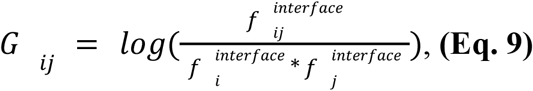

which is the familiar form pairwise residue potentials normalized by interface frequency.(Moont et al., 1999)

### Calculating propensities for specific protein sequences

The stickiness S of a particular protein was calculated as:

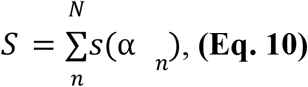

where *a*_*n*_ is the amino acid identity at residue *n* of the client protein with *N* sites, and *s* is the stickiness value of amino acid *a*_*n*_ as obtained from Eq. 1 and available in Supplementary Table 1.

The interaction potential between the client and the droplet is calculated as:

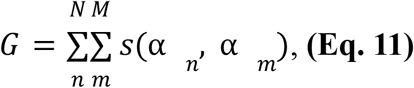

where *a*_*n*_ is the amino acid identity at residue *n* of a client protein with *N* sites, and *a*_*m*_ is the amino acid identity at residue *m* of the partner protein with *M* sites, and *G* is the pairwise potential value. In this equation, we ignore any internal structure in the droplet-client complex and assume that all amino-acids of the client and partner interact with equal probability. This is consistent with observations that interactions within biomolecular condensates are heterogenous and not limited to specific pairwise interactions.(Murthy et al., 2019)

## RESULTS AND DISCUSSION

### A statistical potential unifying residue-solvent and residue-residue interactions

The binding of a protein surface to a partner molecule involves two major energetic components. A first is the desolvation of amino-acids forming the new interface, and a second stems from contacts and non-covalent interactions established across the interface. We first aim to derive statistical potentials enabling a direct comparison of the energetic contribution of both of these components. Such comparison is made possible by calculating the ratios of amino acid surface areas, either (i) between protein surfaces and interfaces to estimate desolvation terms, or (ii) between total interfaces and a sub-part of interfaces composed of a specific amino acid to estimate contact preferences between amino acid residues (Figure 1, Figure 2A).

**Figure 2.**
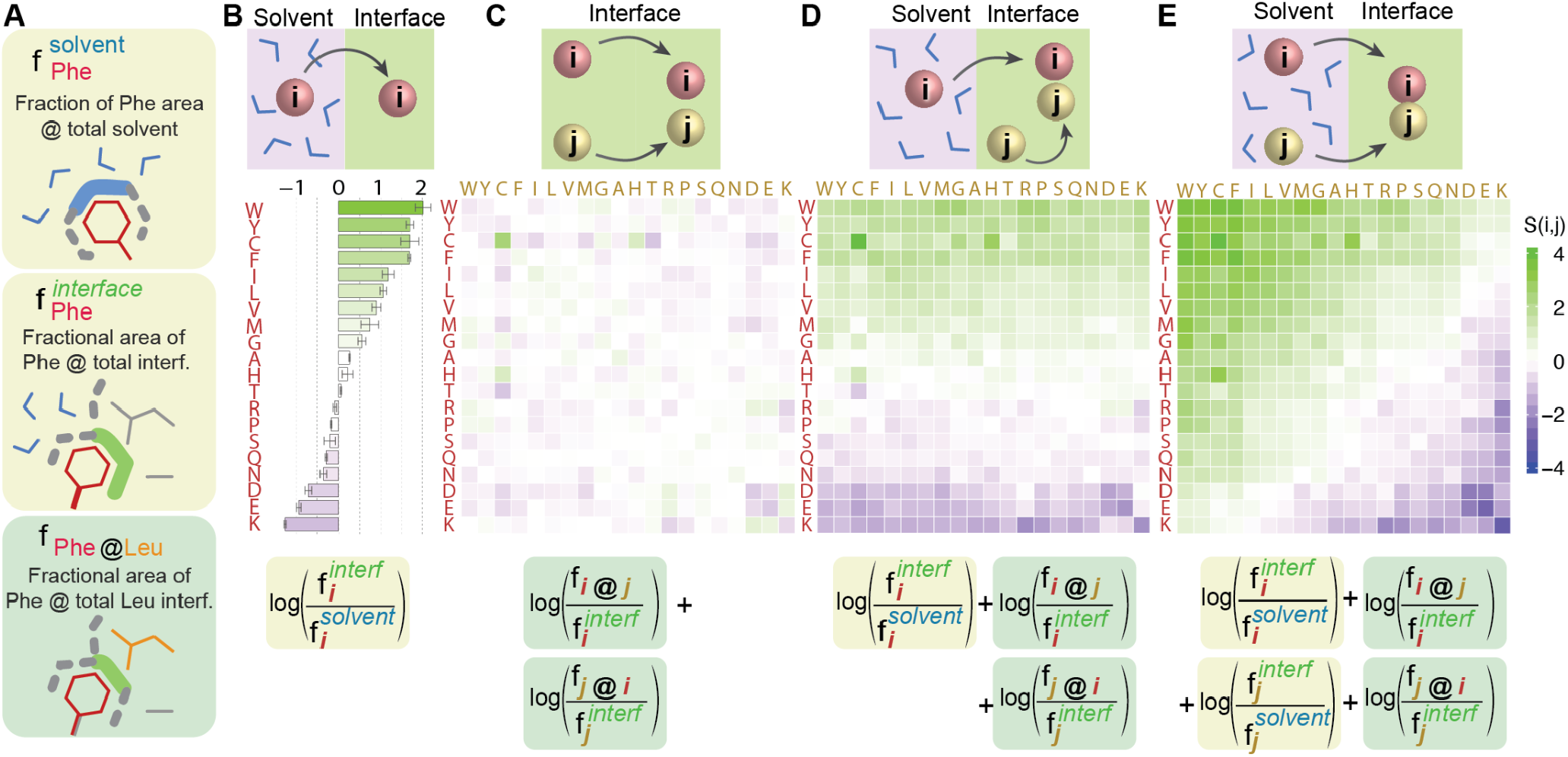
Defining amino-acid interface propensities and interaction propensities based on surface areas. **A**. We calculate three types of surface areas to derive interface propensities and pairwise interaction propensities: (i) The area fraction an amino occupies at solvated surfaces. Phenylalanine, for example, makes up 1.33% of all protein surfaces in our dataset. (ii) The area fraction an amino acid occupies at interfaces. Phenylalanine, for example, makes up 7.19% of all protein interfaces in our dataset. (iii) The area fraction an amino acid makes up at a sub-interface region defined by a particular amino acid. For example, phenylalanine makes up 8.17% of the total leucine interface area. **B**. We estimate the free energy of transfer of amino acids from solvent to interface from the statistics of surface areas contributed to both regions. For example, the interface propensity of phenylalanine is log(0.0719/0.0133) = 1.69. **C**. We estimate the interaction propensity of amino-acids independently of their desolvation component. While the area fraction of phenylalanine at the total interface is 7.19%, it contacts 8.17% of leucine’s interface area, highlighting a representation of this contact that is close to a random expectation: log(0.0817/0.0719)=0.13. **D**. We estimate the interaction propensity of amino-acids with amino-acid *i* being desolvated (red) and amino acid *j* (yellow) being already at the interface. **E**. Interaction propensity that includes the desolvation component for both amino acids *i* and *j*.

Calculating the partitioning of amino acid contact areas at interfaces versus at the solvent yields a 1D stickiness scale (Figure 2B, Equation 1). Consistent with previous interface propensity scales, lysine has the lowest propensity followed by negatively charged and polar amino acids, while aromatic and hydrophobic amino acids have the highest. Also consistent with previous scales and observations, arginine is significantly more sticky than lysine despite being highly hydrophilic owing to its increased ability to establish various contact with partner amino acids.(Conte et al., 1999; Dai et al., 2016; Levy et al., 2012; Yan et al., 2008)

Pairwise interactions are inferred from the area of contact between a pair of amino acids normalized by the area of those amino acids at interfaces.(Figure 2C, Equation 6) This is akin to a pairwise interaction propensity scale. As expected, oppositely charged residues exhibit the highest interaction propensities. Interestingly, we find that a majority of non-charged amino-acids (A, C, F, G, H, I, L, N, S, T, V) exhibit favorable self-interactions whereby the mean potential is larger than the standard deviation, perhaps reflecting on the natural tendency of proteins for self-interaction and self-aggregation (Lukatsky et al., 2007; Wright et al., 2005). Of all these amino acids, cysteine shows the most favorable self-interaction potential. Importantly, this observation is not caused by symmetric homodimers as our dataset is composed of heterodimers exclusively (Methods). Important also, unlike the 1D stickiness, these pairwise interactions do not include desolvation and only relate to contact preference *within* an already formed interface.

Decomposing the interaction propensity into separate energetic terms enables the consideration of the asymmetric interaction between two amino acid residues across a phase boundary. If we consider an environment of reduced hydration, such as the protein dense phase of a phase-separated system, we can add the desolvation term to only the amino acid residues of the client protein, while considering the desolvation states of the scaffold residues as remaining unchanged (Figure 2D).

Finally, we can add the desolvation term to both amino acids entering in contact, resulting in another familiar form of an interaction matrix that considers desolvation (Figure 2E). The interaction energies of oppositely charged residues were among the most favourable in the pairwise interaction matrix that does not consider solvation. Interestingly, these favorable interactions are now offset by the unfavorable desolvation energies. Thus, although most pairwise residue potentials would classify a Lys-Asp interaction as favorable, our potential describes it as unfavorable when considering both interaction energies and desolvation effects. This is reflective of the fact that lysine prefers to be in contact with the solvent, regardless of the existence of favorable electrostatic interactions with glutamate. Similarly Arg-Asp interactions are unfavorable, albeit close to a neutral (zero) value owing to the higher interface propensity of arginine. We see an opposite trend with tyrosine and tryptophan, which show favorable interactions with all amino acids due to a highly positive desolvation term.

Overall, this matrix of pairwise interaction potentials recapitulates the early observation that desolvation is driving complex formation, whereas electrostatic interactions tune interaction specificity. (Chothia and Janin, 1975)

### Analysis of client partitioning within FG domains of the NPC

The NPC is a large protein complex regulating the transport of biomolecules across the nuclear membrane. A hallmark of the NPC are long disordered regions rich in phenylalanine and glycine (FG-domains) that fill up the central cavity and form a gel-like structure thought to phase-separate.(Celetti et al., 2020; Schmidt and Görlich, 2015; Wente et al., 1992) An important step in the transport of cargo across the nuclear pore is the dissolution of the cargo within the phase-separated FG domains, which is dependent on the cargo’s composition. Frey *et al*. characterized the partitioning of protein cargo within liquid protein droplets composed of FG-domain containing sequences. They found that the partitioning of protein cargo coincided with the passage of cargo across the nuclear pore complex. (Frey et al., 2018) GFP variants that differed only in the identity of a single amino acid type at eight different positions on the protein surface were synthesized, and the partition coefficients of each variant between the dilute phase and the FG domain phase were measured (Figure 3A).

**Figure 3.**
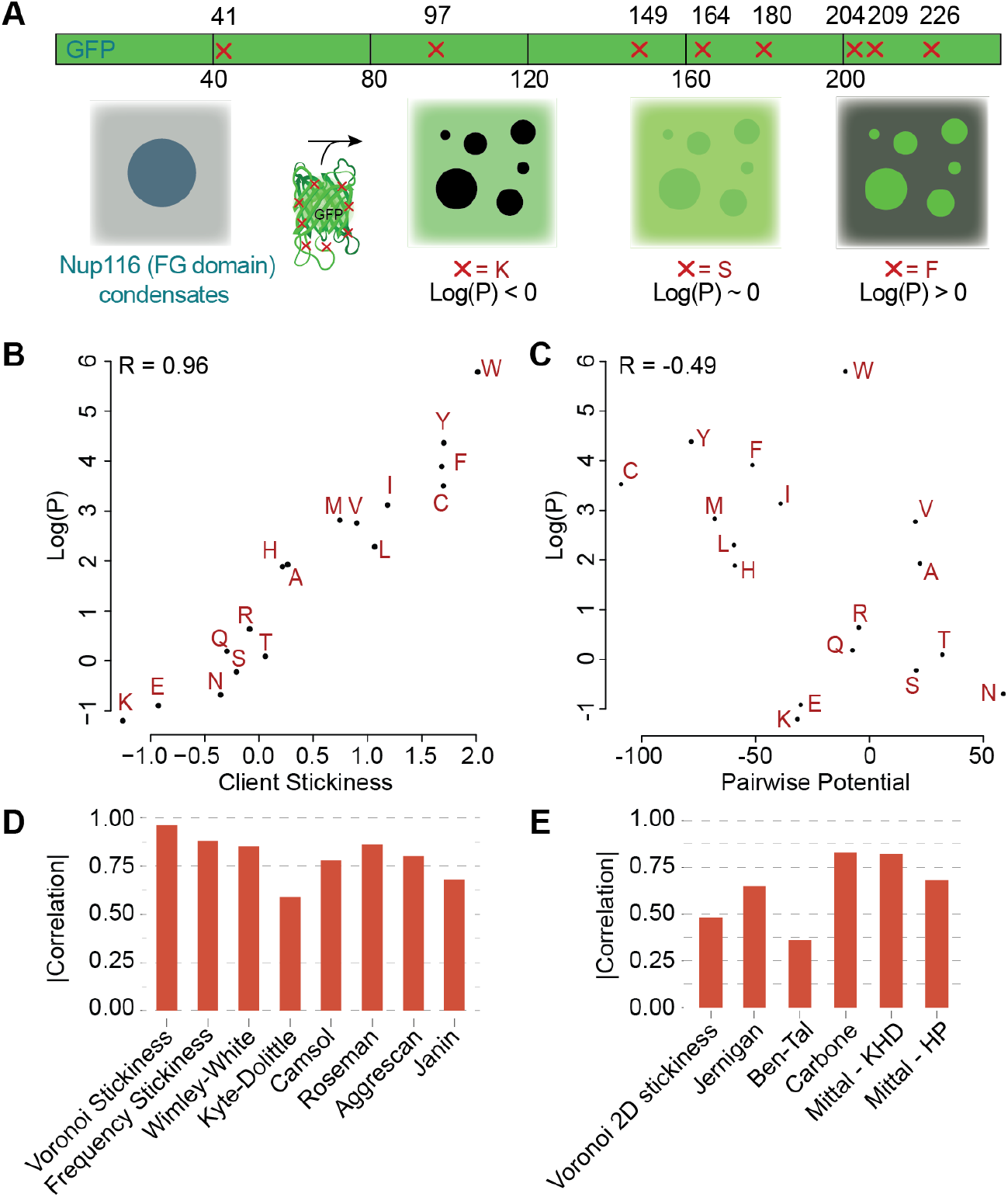
Residue desolvation predicts the dissolution of a protein cargo into FG-rich condensates better than residue-residue interactions. **A**. Frey et al. (Frey et al., 2018) generated GFP variants harboring eight mutations at their surface. In each variant, all eight mutations were to the same amino acid. They measured the partition coefficient (log(P)) of each variant between bulk and condensates made of a FG-rich sequence from Nup116. **B**. The stickiness scale derived in this work recapitulates the observed partition coefficients well. **C**. The residue-residue interaction energies derived in this work do not explain the partition coefficients observed, indicating the desolvation energy is driving the dissolution of cargo into these condensates. The Gle2-binding (GLEB) domain was not included in calculating the interaction potential. **D**. We assessed several hydropathy and solubility scales for their ability to recapitulate the observed partition coefficients. **E**. We assessed several residue-residue interaction potentials for their ability to recapitulate the observed partition coefficients.

We plotted the log-values of the partition coefficients with the scores calculated with the use of our derived scales. The derived residue propensity values for each amino acid is highly correlated with the partition coefficient of each of the variants in systems composed of phase-separated NUP droplets (*R* = 0.96). This high correlation implies that the desolvation energy of the client protein is the main driver of the partitioning between the two phases. We compared our results to those obtained with other propensity scales and pairwise residue potentials. We first analysed a set of hydrophobicity scales. These were Wimley-White,(Wimley and White, 1996) Kyte-Doolittle,(Kyte and Doolittle, 1982) CamSol,(Sormanni et al., 2015) Roseman,(Roseman, 1988) Janin,(Janin, 1979) and Aggrescan,(Conchillo-Solé et al., 2007). As can be seen in Figure 2C, our Voronoi-based residue interface propensity scale is able to capture the partition of proteins to a better degree.

We now consider pairwise interaction energies (Figure 2C) between the amino acid of interest and amino acids in the sequence of NUP116, which are summed as described in Eq. 11. The total interaction energies so obtained for the different variants correlate negatively with the partition coefficients (*R =* -0.49). This indicates that the contribution of specific pairwise interactions between the client protein and the scaffold is negligible in establishing the partitioning. We then compared this correlation value to those derived in the same manner (Eq. 11), based on several prominent pairwise propensity scales. These include classical Jernigan potential(Miyazawa and Jernigan, 1985) and the Glaser scale from the Ben-Tal group.(Glaser et al., 2001) The Mittal group specifically developed one-dimensional scales for modeling phase separation by protein disordered regions, where pair interactions were calculated as the average single amino acid values,(Dignon et al., 2018)he Carbone group developed a scale that combines interaction potential with interface propensity, thus incorporating desolvation energies into each term.(Nadalin et al., 2018) Although the pair propensities in Glaser scale reflect the tendency for hydrophobic amino acids to interact at protein interfaces, this scale was designed to capture amino acid interaction energies without desolvation contributions. This lack of an explicit hydrophobic term is reflected in the reduced performance of this scale compared to scales that account for desolvation, further indicating the desolvation energy is the main driver of the partitioning.

### Analysis of client partitioning within FUS droplets

In a different system, Wang *et al*. investigated the partitioning of proteins within protein liquid droplets made from the protein Fused in Sarcoma (FUS).(Wang et al., 2018) In these experiments, FUS is fused to a SNAP-tag conjugated to a red fluorescent dye and upon phase separation the dense phase is visible as a red-fluorescent droplet. After mixing with various GFP-fused client proteins, a partition coefficient is calculated from the green fluorescence intensity inside versus outside of droplets (Figure 4A). As a first approximation, the authors described the dependence of the partition coefficients on the number of arginine and tyrosine residues in the disordered regions of the client proteins. Indeed, tyrosine and arginine residues have been observed to be overrepresented in certain proteins prone to undergo phase separation, consistent with the observation that cation-П interactions can drive condensate formation.(Nott et al., 2015; Qamar et al., 2018) As these types of interactions are known to be important contributors to protein folding, Das *et. al*. used coarse-grained simulations to find that empirically derived potentials trained on folded proteins can correctly capture cation-П interaction propensities in phase-separating systems.(Das et al., 2020) We also observed a Pearson’s correlation between client sequence length of disordered regions and the log of the partition coefficients of 0.71. This can be seen as analogous to buried interface surface area in protein complexes. Interestingly, the sole number of tyrosine residues in client proteins was a strong predictor of the partition coefficient (R=0.95), as were the tyrosine plus arginine counts (R=0.86).

**Figure 4.**
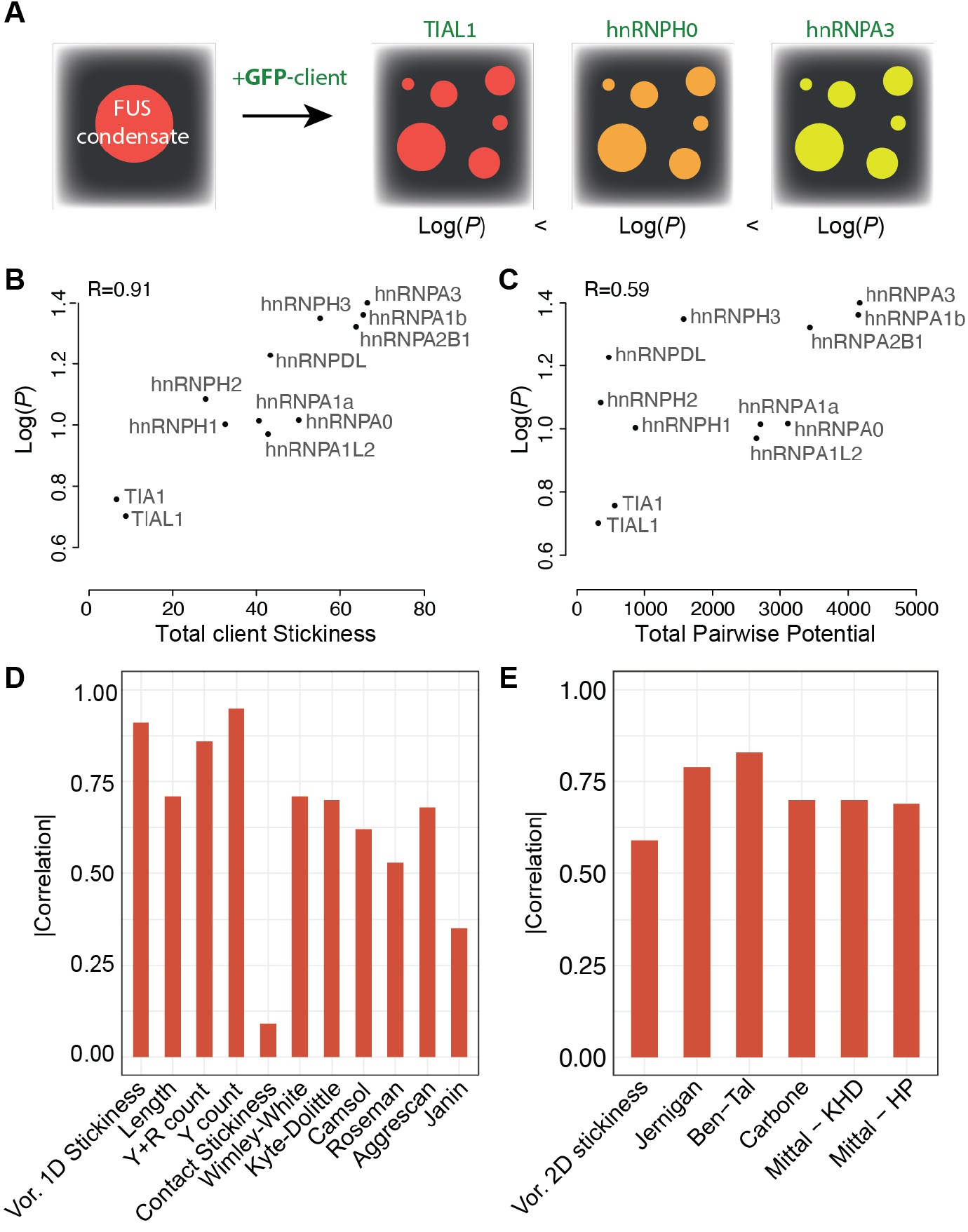
Comparing desolvation energy and residue-residue contact energies in their ability to predict client recruitment into FUS condensates. **A**. FUS condensates exhibiting red fluorescence are mixed with various clients and a partition coefficient is measured for each client. **B**. Total stickiness of each client (x-axis) as a function of the partition coefficient. **C**. Total contact preference potential between each client and FUS. **D**. Correlation between several client’s sequence features and their partition coefficient. Certain features are derived from the sequence directly (e.g., length, Y count) while others correspond to a summed potential based on the same hydropathy and solubility scales used previously. **E**. Correlation between *log(P)* and client-FUS contacts calculated based on several contact potentials.

To shed more light on the sequence dependence of the client in the partitioning within FUS droplets, we correlated the total stickiness of the disordered regions of client proteins to the log values of the measured partition coefficients. We also correlated the stickiness of each client to FUS by summing over all pairwise interactions between client and host. As we saw before, the total stickiness reflects the desolvation energy of the client, whereas pairwise interactions as shown in Figure 2C and used in Eq. 11 reflect residue-residue contact preferences without desolvation being factored in the potential. We observed a high correlation between the total stickiness of the client protein and the log of the partition coefficient (R=0.91, Figure 4B). In contrast, the contact potential gave a substantially lower correlation (R=0.59, Figure 4C). While all other pairwise potentials tested performed better than our Voronoi-derived scale, none of the potentials approached the performance of metrics that only considered client sequence properties. In fact, most of the scales used were predictive to the same degree as client sequence length alone. A simple count of tyrosine residues in the sequence proved to be the best predictor, so that the reason the Glaser scale outperforms in the pairwise case is likely due to the fact that it exhibits strong preferences for interactions involving bulky aromatic amino acids.

A striking result of our analysis was the substantial underperformance of the stickiness scale calculated from interface contact counts. A comparison of the Voronoi 1D stickiness and contact stickiness scales reveals that Gly stickiness differs between the two scales, where Gly is determined to be unfavorable at interfaces when contact counts are used. Given that intrinsically disordered protein segments are rich in glycine residues that confer flexibility, the stickiness value of Gly is a major component of the total stickiness value of each client protein. Replacing the value of Gly stickiness in the contact stickiness scale (−0.1771) by the value in the Voronoi stickiness scale (0.57) results in a correlation of 0.85 between the total stickiness of the client proteins and the log of partition coefficients. This illustrates the importance of considering the physico-chemical properties of all residues in phase-separated systems.

## Discussion and conclusions

*In vitro* condensate formation results in a non-homogenous solution composed of two distinct chemical environments. Additional molecules are distributed throughout this heterogeneous environment in such a way that minimizes the free energy of the system. Entropic and enthalpic factors both contribute to the energy minimization, and our results indicate that entropy changes due to water organization around hydrophobic surfaces of client molecules play a significant role. This result may appear surprising because condensates are expected to remain highly hydrated. For example, the water content in a phase-separating system composed of ε-poly-L-lysine (εPL) and hyaluronic acid (HA) was measured at 81.18%.(Park et al., 2020) Indeed, even in protein crystals, a significant volume is occupied by bulk water.(Matthews, 1968) In that respect, the desolvation of the client molecules does not only require the client’s presence in the dense phase, but also requires interactions with the species forming the condensate. However, our results imply that these interactions are non-specific and that client partitioning is driven by the increase in the entropy of water within the droplet that occurs as a result.

We reasoned that the interface of protein complexes would provide a reasonable proxy for estimating the chemical environment in the dense phase of biomolecular condensates, while the protein surface would provide the same for the dilute environment. In doing so, we expected that using data extracted from structured protein-protein interfaces would not be an impediment for using such potentials to describe highly dynamic systems, as the underlying physical nature of amino acid desolvation and amino acid pair interactions is the same. We used Voronoi surface area as a quantity that could be used to directly compare amino acid interface and the surface occupancy, and developed a novel interface propensity scale and a pairwise residue propensity score. Uniquely, the two potentials are orthogonal to each other and they can be combined as they are derived from the same information type. That is, amino acid surface areas and contact areas.

We find that the interface propensity scores provide good correlation when used to examine the partitioning of client proteins in phase separated systems. When we calculated pairwise interaction energies, however, we did not observe the same degree of correlation. This suggests that the main driver of client partitioning is the desolvation energy of going from a dilute environment to a protein dense environment. The reduced performance of the pairwise interaction scale can be partially explained by our lack of explicit sampling of client-droplet configurations, thus ignoring the fact that some specific amino acid pair interactions could be dominant. However, the fact that client partitioning correlates to such a large degree with interface propensity implies that the specificity of these interactions have a comparatively small contribution in the free energy of interactions. Interestingly, this observation indicates that specific interactions between purely disordered sequences may not be sufficient to build selectivity into client partitioning in liquid protein systems. And consequently, additional recognition features are likely required for selectivity. This notion is in contrast to the observations made regarding the sequence determinants of phase-separation propensities of scaffold proteins, which are not adequately accounted for with desolvation energies alone, (Dignon et al., 2020) and where composition as well as sequence patterns play an important role.(Borcherds et al., 2021; Hazra and Levy, 2020; Martin et al., 2020)

Tools for predicting the distribution of species in the complex environment of the cell are crucial for understanding cellular organization as well as pharmacokinetic behaviour of therapeutic drugs. As the number of newly discovered membraneless organelles, as well as the understanding of the biological processes that are governed by phase-separation, the development of potentials that can closely reflect the partitioning of proteins within liquid protein droplets will enable the design of synthetic systems that could be used for the regulation of cellular processes.

## Supporting information

2D plus 1D Symmetric Scale

2d plus 1D Scale

2D Scale SD

2D Scale

1D Scale SD

1D Scale

List of PDBs

## Notes

### Competing Interest Statement

The authors have declared no competing interest.

## References

Alberts, B. (2017). Molecular Biology of the Cell (Garland Science).

Banani, S.F., Lee, H.O., Hyman, A.A., and Rosen, M.K. (2017). Biomolecular condensates: organizers of cellular biochemistry. Nat. Rev. Mol. Cell Biol. 18, 285–298.

Borcherds, W., Bremer, A., Borgia, M.B., and Mittag, T. (2021). How do intrinsically disordered protein regions encode a driving force for liquid-liquid phase separation? Curr. Opin. Struct. Biol. 67, 41–50.

Brangwynne, C.P., Eckmann, C.R., Courson, D.S., Rybarska, A., Hoege, C., Gharakhani, J., Jülicher, F., and Hyman, A.A. (2009). Germline P granules are liquid droplets that localize by controlled dissolution/condensation. Science 324, 1729–1732.

Brangwynne, C.P., Mitchison, T.J., and Hyman, A.A. (2011). Active liquid-like behavior of nucleoli determines their size and shape in Xenopus laevis oocytes. Proc. Natl. Acad. Sci. U. S. A. 108, 4334–4339.

Celetti, G., Paci, G., Caria, J., VanDelinder, V., Bachand, G., and Lemke, E.A. (2020). The liquid state of FG-nucleoporins mimics permeability barrier properties of nuclear pore complexes. J. Cell Biol. 219.

Chothia, C., and Janin, J. (1975). Principles of protein–protein recognition. Nature 256, 705.

Conchillo-Solé, O., de Groot, N.S., Avilés, F.X., Vendrell, J., Daura, X., and Ventura, S. (2007). AGGRESCAN: a server for the prediction and evaluation of “hot spots” of aggregation in polypeptides. BMC Bioinformatics 8, 65.

Conte, L.L., Chothia, C., and Janin, J. (1999). The atomic structure of protein-protein recognition sites1. J. Mol. Biol. 285, 2177–2198.

Dai, W., Wu, A., Ma, L., Li, Y.-X., Jiang, T., and Li, Y.-Y. (2016). A novel index of protein-protein interface propensity improves interface residue recognition. BMC Syst. Biol. 10, 112.

Das, R.K., and Pappu, R.V. (2013). Conformations of intrinsically disordered proteins are influenced by linear sequence distributions of oppositely charged residues. Proc. Natl. Acad. Sci. U. S. A. 110, 13392–13397.

Das, S., Lin, Y.-H., Vernon, R.M., Forman-Kay, J.D., and Chan, H.S. (2020). Comparative roles of charge, π, and hydrophobic interactions in sequence-dependent phase separation of intrinsically disordered proteins. Proc. Natl. Acad. Sci. U. S. A. 117, 28795–28805.

Dey, S., and Levy, E.D. (2018). Inferring and Using Protein Quaternary Structure Information from Crystallographic Data. In Protein Complex Assembly: Methods and Protocols, J.A. Marsh, ed. (New York, NY: Springer New York), pp. 357–375.

Dey, S., Ritchie, D.W., and Levy, E.D. (2018). PDB-wide identification of biological assemblies from conserved quaternary structure geometry. Nat. Methods 15, 67–72.

Dignon, G.L., Zheng, W., Kim, Y.C., Best, R.B., and Mittal, J. (2018). Sequence determinants of protein phase behavior from a coarse-grained model. PLoS Comput. Biol. 14, e1005941.

Dignon, G.L., Best, R.B., and Mittal, J. (2020). Biomolecular Phase Separation: From Molecular Driving Forces to Macroscopic Properties. Annu. Rev. Phys. Chem. 71, 53–75.

Ditlev, J.A., Case, L.B., and Rosen, M.K. (2018). Who’s In and Who’s Out—Compositional Control of Biomolecular Condensates. Journal of Molecular Biology 430, 4666–4684.

Frey, S., Rees, R., Schünemann, J., Ng, S.C., Fünfgeld, K., Huyton, T., and Görlich, D. (2018). Surface Properties Determining Passage Rates of Proteins through Nuclear Pores. Cell 174, 202–217.e9.

Fuxreiter, M., and Vendruscolo, M. (2021). Generic nature of the condensed states of proteins. Nat. Cell Biol. 23, 587–594.

Ge, X., Conley, A.J., Brandle, J.E., Truant, R., and Filipe, C.D.M. (2009). In vivo formation of protein based aqueous microcompartments. J. Am. Chem. Soc. 131, 9094–9099.

Ghosh, A., and Mazarakos, K. (2019). Three archetypical classes of macromolecular regulators of protein liquid–liquid phase separation. Proceedings of the.

Glaser, F., Steinberg, D.M., Vakser, I.A., and Ben-Tal, N. (2001). Residue frequencies and pairing preferences at protein-protein interfaces. Proteins 43, 89–102.

Hardenberg, M., Horvath, A., Ambrus, V., Fuxreiter, M., and Vendruscolo, M. (2020). Widespread occurrence of the droplet state of proteins in the human proteome. Proc. Natl. Acad. Sci. U. S. A. 117, 33254–33262.

Hazra, M.K., and Levy, Y. (2020). Charge pattern affects the structure and dynamics of polyampholyte condensates. Phys. Chem. Chem. Phys. 22, 19368–19375.

Howard, M. (2012). How to build a robust intracellular concentration gradient. Trends Cell Biol. 22, 311–317.

Hülsmann, B.B., Labokha, A.A., and Görlich, D. (2012). The permeability of reconstituted nuclear pores provides direct evidence for the selective phase model. Cell 150, 738–751.

Janin, J. (1979). Surface and inside volumes in globular proteins. Nature 277, 491–492.

Jones, S., and Thornton, J.M. (1996). Principles of protein-protein interactions. Proc. Natl. Acad. Sci. U. S. A. 93, 13–20.

Kato, M., Han, T.W., Xie, S., Shi, K., Du, X., Wu, L.C., Mirzaei, H., Goldsmith, E.J., Longgood, J., Pei, J., et al. (2012). Cell-free formation of RNA granules: low complexity sequence domains form dynamic fibers within hydrogels. Cell 149, 753–767.

Kyte, J., and Doolittle, R.F. (1982). A simple method for displaying the hydropathic character of a protein. J. Mol. Biol. 157, 105–132.

Levy, E.D., Pereira-Leal, J.B., Chothia, C., and Teichmann, S.A. (2006). 3D complex: a structural classification of protein complexes. PLoS Comput. Biol. 2, e155.

Levy, E.D., De, S., and Teichmann, S.A. (2012). Cellular crowding imposes global constraints on the chemistry and evolution of proteomes. Proc. Natl. Acad. Sci. U. S. A. 109, 20461–20466.

Li, P., Banjade, S., Cheng, H.-C., Kim, S., Chen, B., Guo, L., Llaguno, M., Hollingsworth, J.V., King, D.S., Banani, S.F., et al. (2012). Phase transitions in the assembly of multivalent signalling proteins. Nature 483, 336–340.

Lin, Y., Protter, D.S.W., Rosen, M.K., and Parker, R. (2015). Formation and Maturation of Phase-Separated Liquid Droplets by RNA-Binding Proteins. Mol. Cell 60, 208–219.

Lin, Y., Currie, S.L., and Rosen, M.K. (2017). Intrinsically disordered sequences enable modulation of protein phase separation through distributed tyrosine motifs. J. Biol. Chem. 292, 19110–19120.

Lin, Y.-H., Forman-Kay, J.D., and Chan, H.S. (2016). Sequence-Specific Polyampholyte Phase Separation in Membraneless Organelles. Phys. Rev. Lett. 117, 178101.

Lukatsky, D.B., Shakhnovich, B.E., Mintseris, J., and Shakhnovich, E.I. (2007). Structural similarity enhances interaction propensity of proteins. J. Mol. Biol. 365, 1596–1606.

Martin, E.W., Holehouse, A.S., Peran, I., Farag, M., Jeremias Incicco, J., Bremer, A., Grace, C.R., Soranno, A., Pappu, R.V., and Mittag, T. (2020). Valence and patterning of aromatic residues determine the phase behavior of prion-like domains. Science 367, 694–699.

Matthews, B.W. (1968). Solvent content of protein crystals. J. Mol. Biol. 33, 491–497.

Miyazawa, S., and Jernigan, R.L. (1985). Estimation of effective interresidue contact energies from protein crystal structures: quasi-chemical approximation. Macromolecules 18, 534–552.

Moont, G., Gabb, H.A., and Sternberg, M.J. (1999). Use of pair potentials across protein interfaces in screening predicted docked complexes. Proteins 35, 364–373.

Murthy, A.C., Dignon, G.L., Kan, Y., Zerze, G.H., Parekh, S.H., Mittal, J., and Fawzi, N.L. (2019). Molecular interactions underlying liquid-liquid phase separation of the FUS low-complexity domain. Nat. Struct. Mol. Biol. 26, 637–648.

Nadalin, F., Carbone, A., and Valencia, A. (2018). Protein–protein interaction specificity is captured by contact preferences and interface composition. Bioinformatics 34, 459–468.

Nott, T.J., Petsalaki, E., Farber, P., Jervis, D., Fussner, E., Plochowietz, A., Craggs, T.D., Bazett-Jones, D.P., Pawson, T., Forman-Kay, J.D., et al. (2015). Phase transition of a disordered nuage protein generates environmentally responsive membraneless organelles. Mol. Cell 57, 936–947.

Nott, T.J., Craggs, T.D., and Baldwin, A.J. (2016). Membraneless organelles can melt nucleic acid duplexes and act as biomolecular filters. Nat. Chem. 8, 569–575.

Olechnovič, K., and Venclovas, C. (2014). The use of interatomic contact areas to quantify discrepancies between RNA 3D models and reference structures. Nucleic Acids Res. 42, 5407–5415.

Pak, C.W., Kosno, M., Holehouse, A.S., Padrick, S.B., Mittal, A., Ali, R., Yunus, A.A., Liu, D.R., Pappu, R.V., and Rosen, M.K. (2016). Sequence Determinants of Intracellular Phase Separation by Complex Coacervation of a Disordered Protein. Mol. Cell 63, 72–85.

Panbianco, C., and Gotta, M. (2011). Coordinating cell polarity with cell division in space and time. Trends Cell Biol. 21, 672–680.

Park, S., Barnes, R., Lin, Y., Jeon, B.-J., Najafi, S., Delaney, K.T., Fredrickson, G.H., Shea, J.-E., Hwang, D.S., and Han, S. (2020). Dehydration entropy drives liquid-liquid phase separation by molecular crowding. Communications Chemistry 3, 1–12.

Qamar, S., Wang, G., Randle, S.J., Ruggeri, F.S., Varela, J.A., Lin, J.Q., Phillips, E.C., Miyashita, A., Williams, D., Ströhl, F., et al. (2018). FUS Phase Separation Is Modulated by a Molecular Chaperone and Methylation of Arginine Cation-π Interactions. Cell 173, 720–734.e15.

Qin, S., and Zhou, H.-X. (2016). Fast Method for Computing Chemical Potentials and Liquid–Liquid Phase Equilibria of Macromolecular Solutions. J. Phys. Chem. B 120, 8164–8174.

Regy, R.M., Dignon, G.L., Zheng, W., Kim, Y.C., and Mittal, J. (2020). Sequence dependent phase separation of protein-polynucleotide mixtures elucidated using molecular simulations. Nucleic Acids Res. 48, 12593–12603.

Ribbeck, K., and Görlich, D. (2001). Kinetic analysis of translocation through nuclear pore complexes. EMBO J. 20, 1320–1330.

Richards, F.M. (1974). The interpretation of protein structures: total volume, group volume distributions and packing density. J. Mol. Biol. 82, 1–14.

Roseman, M.A. (1988). Hydrophilicity of polar amino acid side-chains is markedly reduced by flanking peptide bonds. J. Mol. Biol. 200, 513–522.

Sawle, L., and Ghosh, K. (2015). A theoretical method to compute sequence dependent configurational properties in charged polymers and proteins. J. Chem. Phys. 143, 085101.

Schmidt, H.B., and Görlich, D. (2015). Nup98 FG domains from diverse species spontaneously phase-separate into particles with nuclear pore-like permselectivity. Elife 4.

Schuster, B.S., Dignon, G.L., Tang, W.S., Kelley, F.M., Ranganath, A.K., Jahnke, C.N., Simpkins, A.G., Regy, R.M., Hammer, D.A., Good, M.C., et al. (2020). Identifying sequence perturbations to an intrinsically disordered protein that determine its phase-separation behavior. Proceedings of the National Academy of Sciences 117, 11421–11431.

Simm, S., Einloft, J., Mirus, O., and Schleiff, E. (2016). 50 years of amino acid hydrophobicity scales: revisiting the capacity for peptide classification. Biol. Res. 49, 31.

Singh, R.K., Tropsha, A., and Vaisman, I.I. (1996). Delaunay tessellation of proteins: four body nearest-neighbor propensities of amino acid residues. J. Comput. Biol. 3, 213–221.

Sormanni, P., Aprile, F.A., and Vendruscolo, M. (2015). The CamSol method of rational design of protein mutants with enhanced solubility. J. Mol. Biol. 427, 478–490.

Vernon, R.M., and Forman-Kay, J.D. (2019). First-generation predictors of biological protein phase separation. Curr. Opin. Struct. Biol. 58, 88–96.

Vernon, R.M., Chong, P.A., Tsang, B., Kim, T.H., Bah, A., Farber, P., Lin, H., and Forman-Kay, J.D. (2018). Pi-Pi contacts are an overlooked protein feature relevant to phase separation. Elife 7.

Wang, J., Choi, J.-M., Holehouse, A.S., Lee, H.O., Zhang, X., Jahnel, M., Maharana, S., Lemaitre, R., Pozniakovsky, A., Drechsel, D., et al. (2018). A Molecular Grammar Governing the Driving Forces for Phase Separation of Prion-like RNA Binding Proteins. Cell 174, 688–699.e16.

Wente, S.R., Rout, M.P., and Blobel, G. (1992). A new family of yeast nuclear pore complex proteins. J. Cell Biol. 119, 705–723.

Wimley, W.C., and White, S.H. (1996). Experimentally determined hydrophobicity scale for proteins at membrane interfaces. Nat. Struct. Biol. 3, 842–848.

Wright, C.F., Teichmann, S.A., Clarke, J., and Dobson, C.M. (2005). The importance of sequence diversity in the aggregation and evolution of proteins. Nature 438, 878–881.

Yan, C., Wu, F., Jernigan, R.L., Dobbs, D., and Honavar, V. (2008). Characterization of protein-protein interfaces. Protein J. 27, 59–70.

Zaslavsky, B.Y., and Uversky, V.N. (2018). In Aqua Veritas: The Indispensable yet Mostly Ignored Role of Water in Phase Separation and Membrane-less Organelles. Biochemistry 57, 2437–2451.

Zwicker, D., Decker, M., Jaensch, S., Hyman, A.A., and Jülicher, F. (2014). Centrosomes are autocatalytic droplets of pericentriolar material organized by centrioles. Proc. Natl. Acad. Sci. U. S. A. 111, E2636–E2645.

